## Supplementary Information for "DNA 5-methylcytosine detection and methylation phasing using PacBio circular consensus sequencing"

Ni *et al.*

### Supplementary Figures

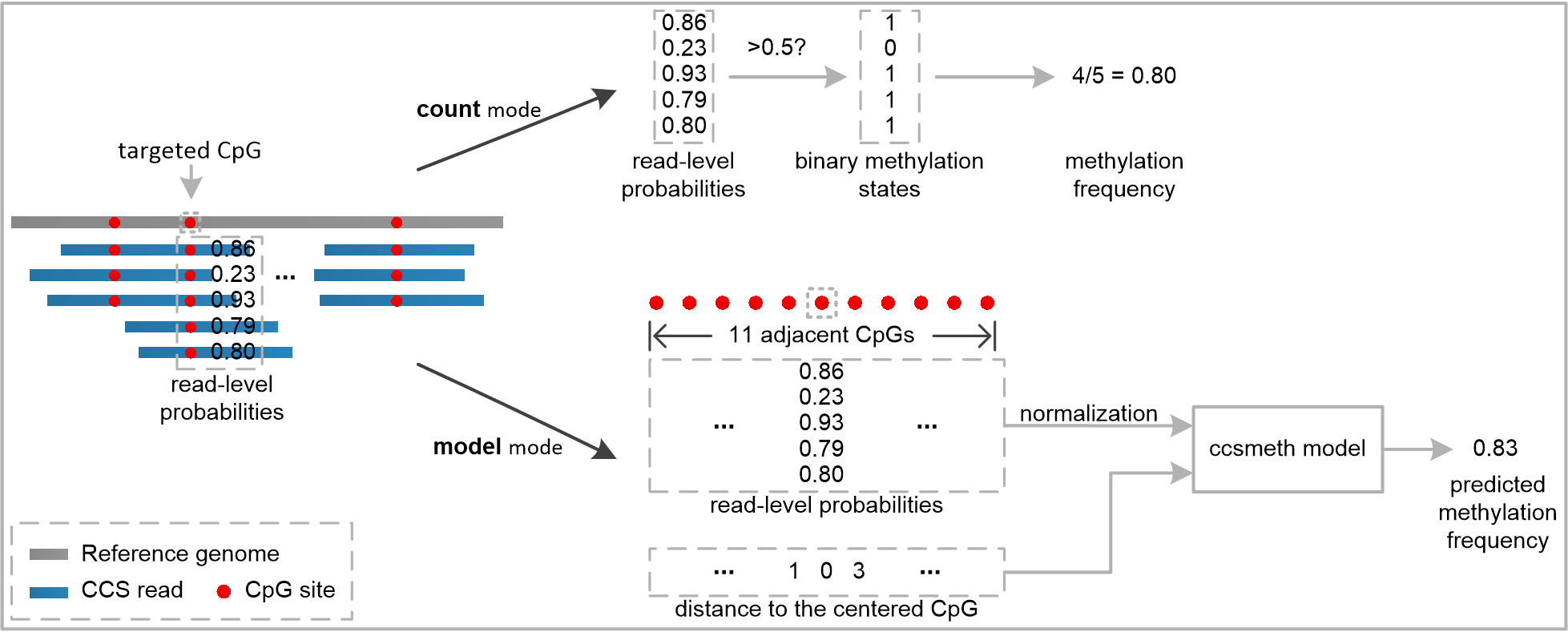

**Supplementary Fig. 1** Illustration of inferring methylation frequency of CpGs at site level using count mode and model mode.

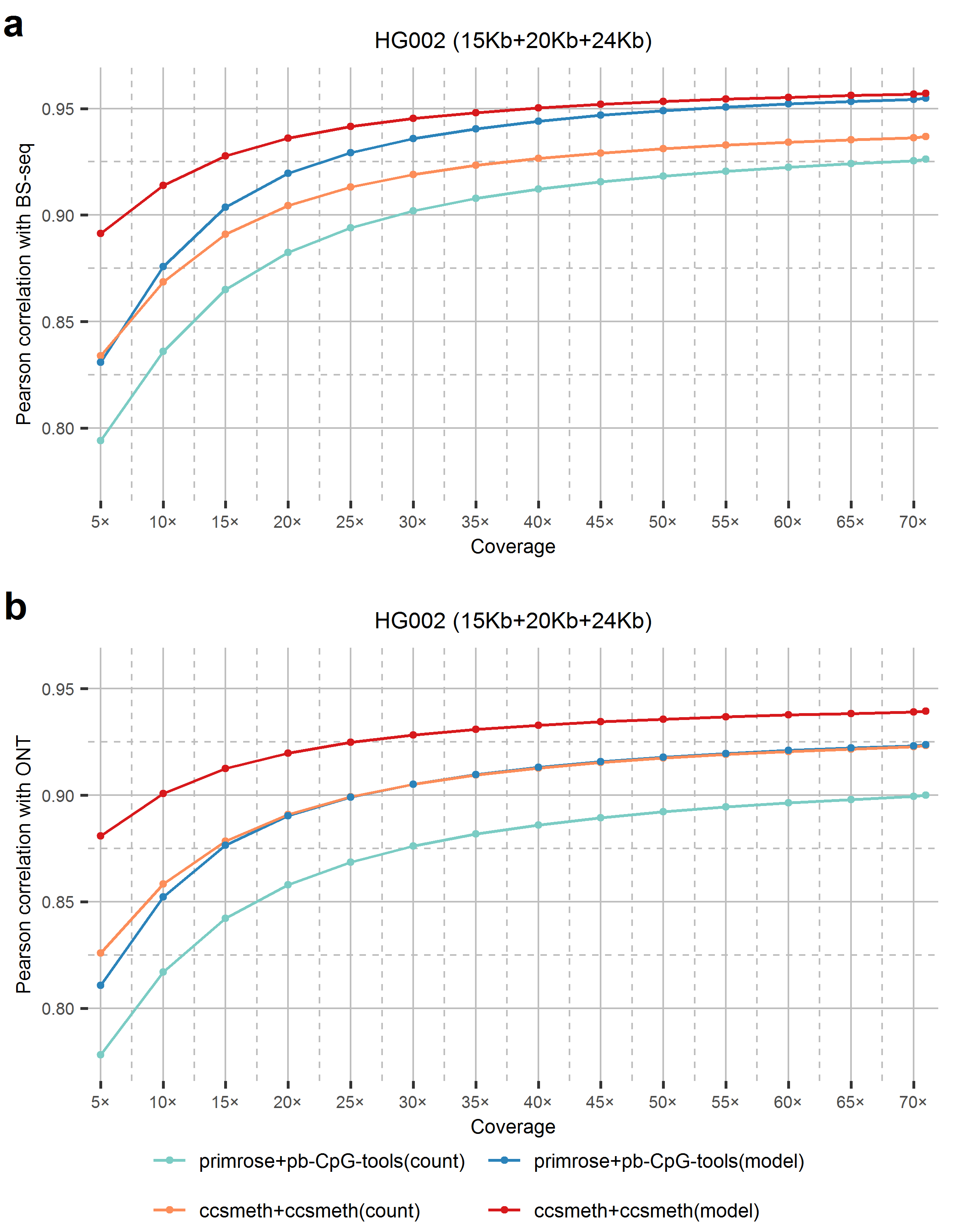

**Supplementary Fig. 2** Comparing ccsmeth and primrose/pb-CpG-tools against BS-seq (**a**) and nanopore sequencing (**b**) under different coverages of HG002 CCS reads (71.0× in total). Values for coverage 5×-70× are the average of 5 repeated tests.

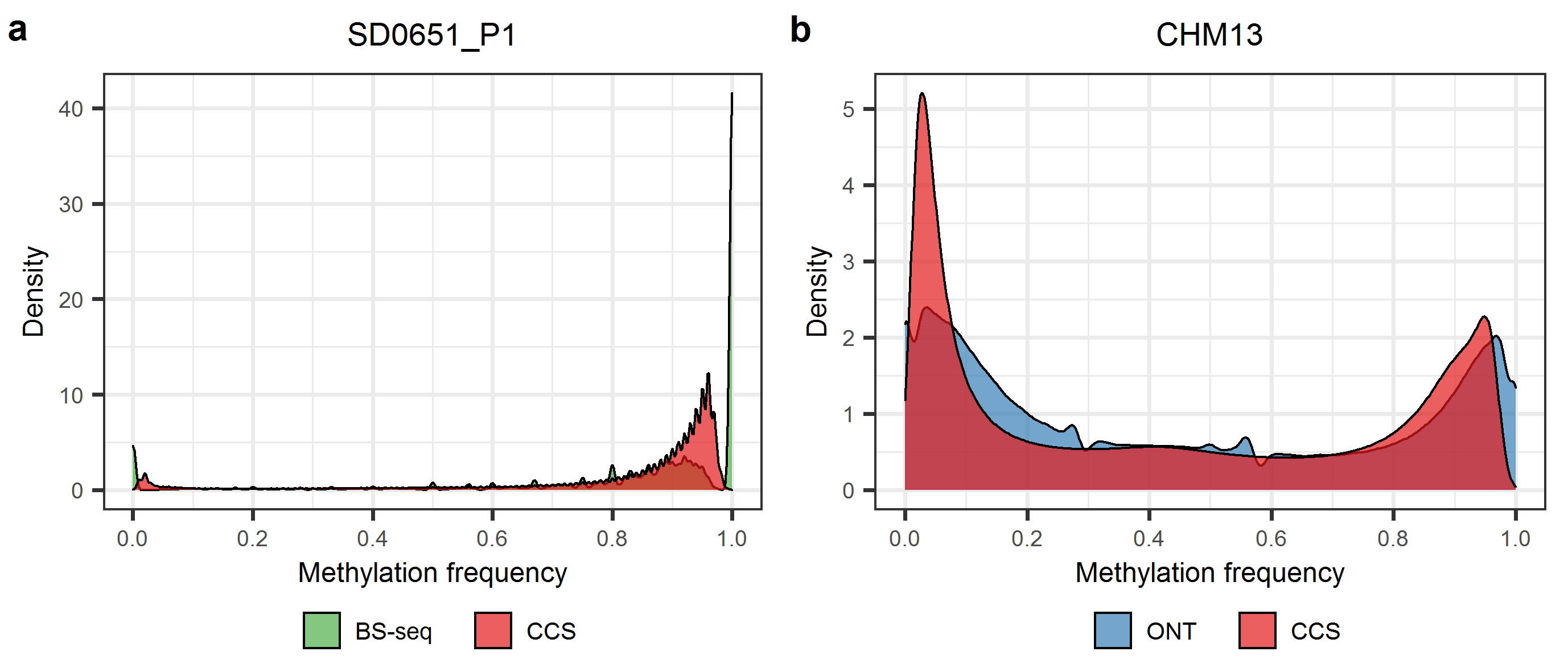

**Supplementary Fig. 3** Methylation frequencies of CpGs in two human samples SD0651_P1 and CHM13. **a** Distribution of methylation frequencies of CpGs in SD0651_P1 detected by BS-seq and CCS (ccsmeth in model mode). **b** Distribution of methylation frequencies of CpGs in CHM13 detected by nanopore sequencing and CCS (ccsmeth in model mode). ONT: nanopore sequencing.

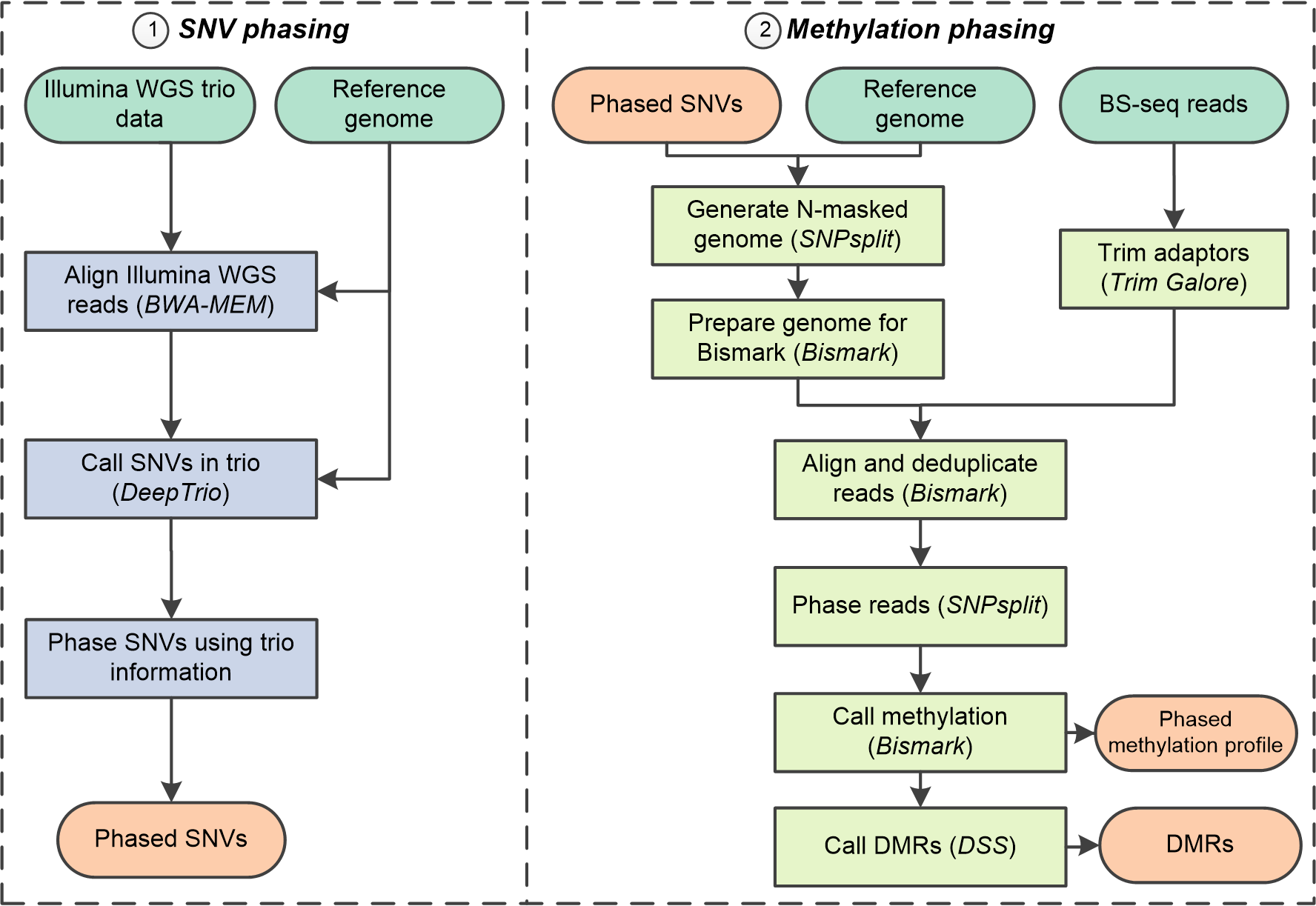

**Supplementary Fig. 4** Pipeline of haplotype-aware methylation calling and allele-specific methylation detection using Illumina whole-genome sequencing (WGS) trio data and BS-seq data. SNVs: single nucleotide variants; DMRs: differentially methylated regions.

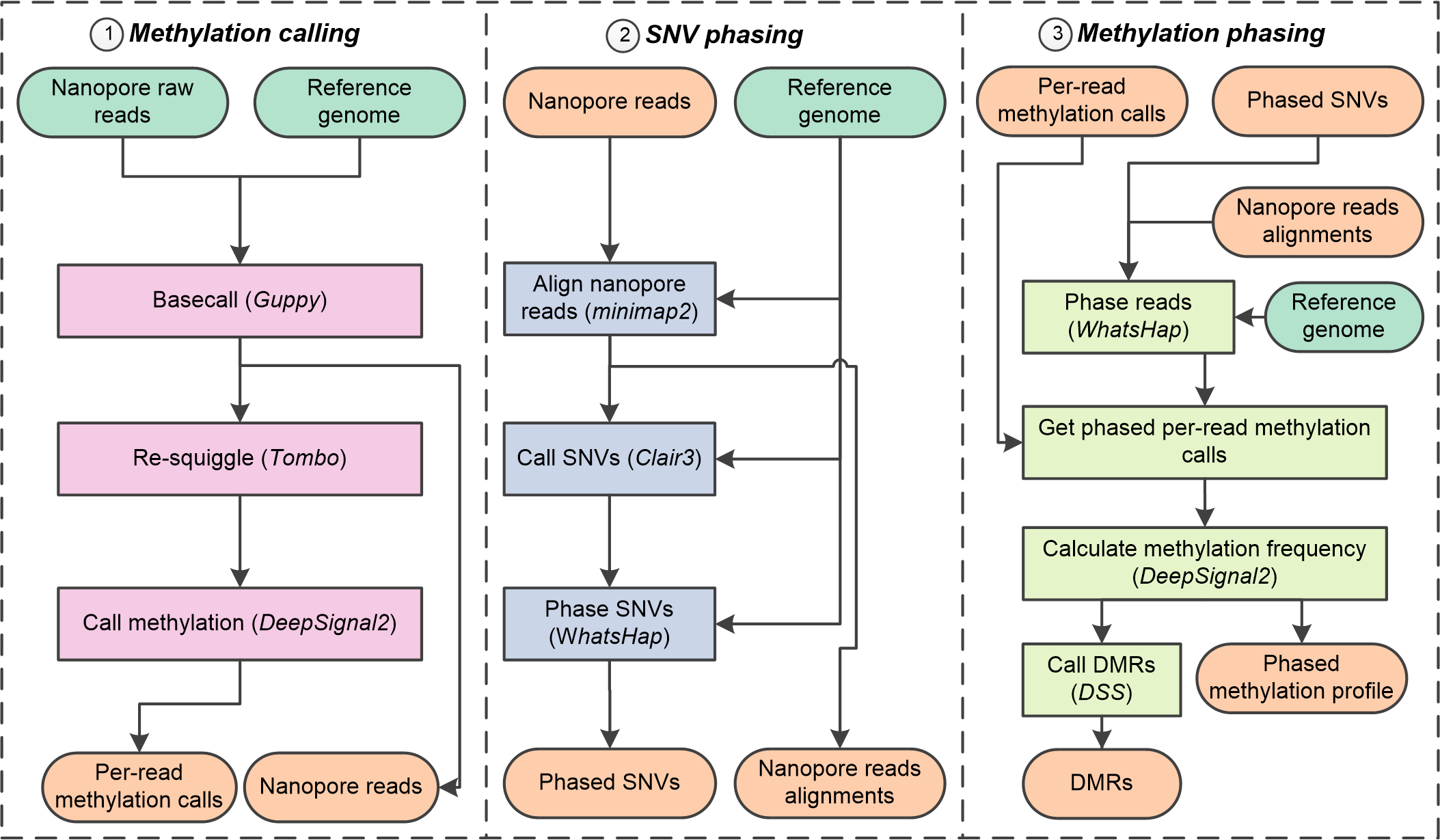

**Supplementary Fig. 5** Pipeline of haplotype-aware methylation calling and allele-specific methylation using nanopore data only. SNVs: single nucleotide variants; DMRs: differentially methylated regions.

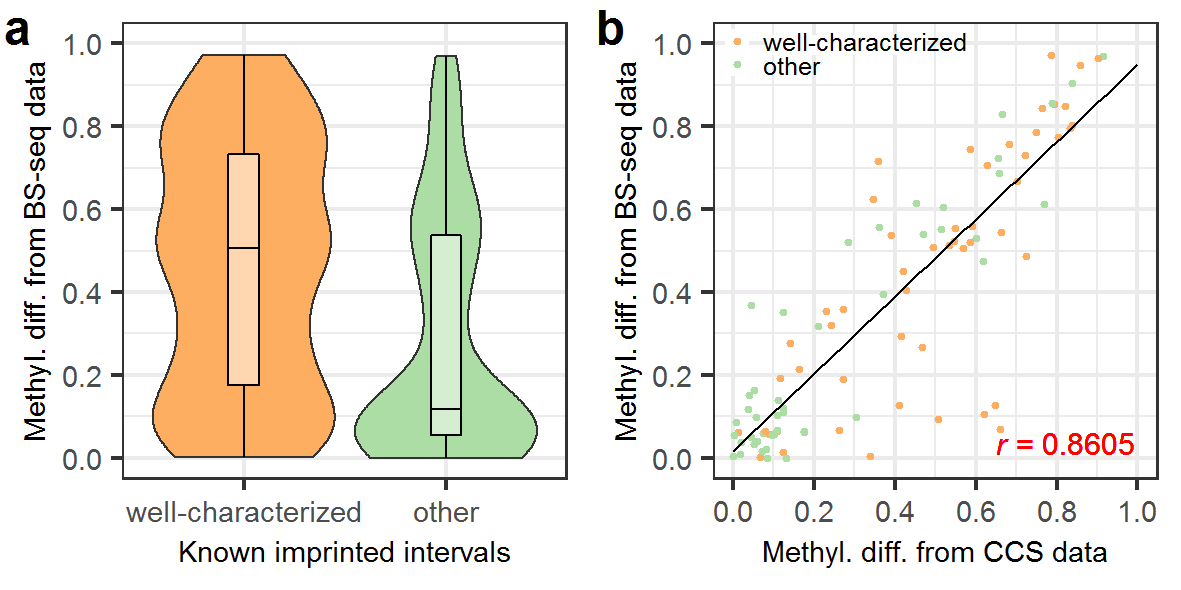

**Supplementary Fig. 6** Methylation differences of known imprinted intervals calculated using BS-seq/CCS data in two haplotypes of HG002 **a** Distribution of methylation differences of known imprinted intervals between two haplotypes of HG002 calculated using BS-seq data. 52 out of 102 well-characterized intervals and 46 out of 102 other intervals which have at least 5 CpGs covered by BS-seq reads in each haplotype are analyzed. The boxes inside the violin plots indicate the 50th percentile (middle line), 25th, and 75th percentile (box). **b** Comparison of methylation differences of known imprinted intervals calculated using CCS and BS-seq data. 97 known imprinted intervals which can be covered by BS-seq and CCS data were analyzed. Methyl. diff.: methylation difference; *r*: Pearson correlation.

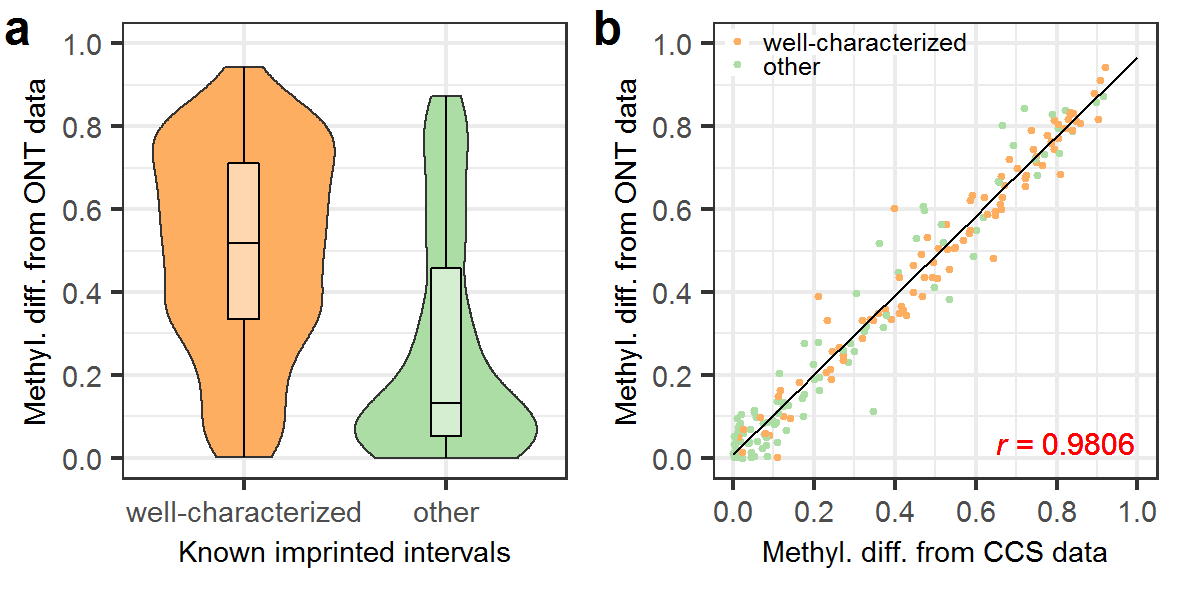

**Supplementary Fig. 7** Methylation differences of known imprinted intervals calculated using nanopore/CCS data in two haplotypes of HG002. **a** Distribution of methylation differences of known imprinted intervals between two haplotypes of HG002 calculated using nanopore data. 98 out of 102 well-characterized intervals and 96 out of 102 other intervals which have at least 5 CpGs covered by nanopore reads in each haplotype are analyzed. The boxes inside the violin plots indicate the 50th percentile (middle line), 25th, and 75th percentile (box). **b** Comparison of methylation differences of known imprinted intervals calculated using CCS and nanopore data. 191 known imprinted intervals which can be covered by nanopore and CCS data were analyzed. Methyl. diff.: methylation difference; ONT: nanopore sequencing; *r*: Pearson correlation.

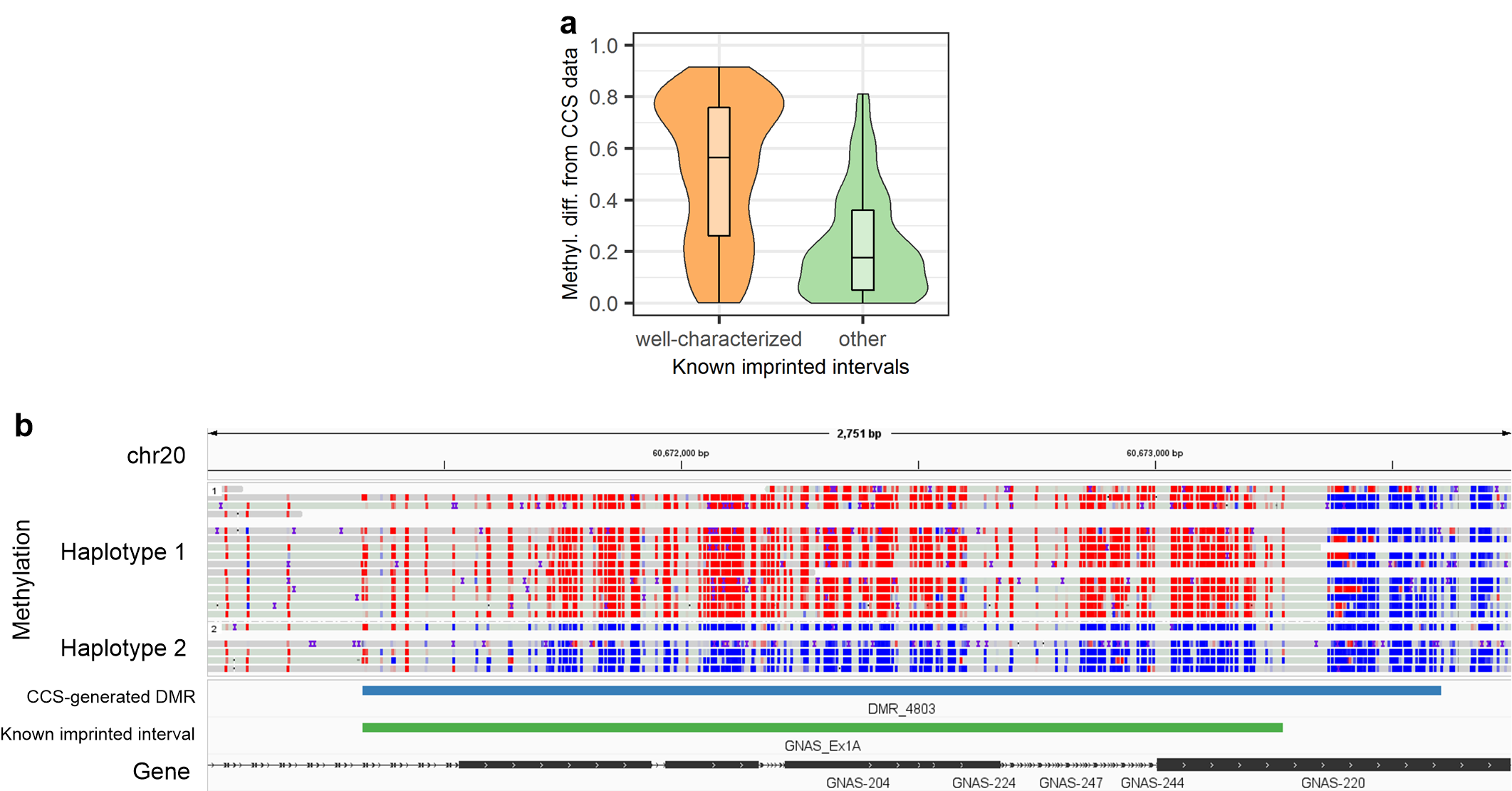

**Supplementary Fig. 8** Methylation phasing of ccsmethphase on SD0651_P1 CCS data. **a** Distribution of methylation differences of known imprinted intervals calculated using CCS data between two haplotypes of SD0651_P1. 93 out of 102 “well-characterized” intervals, and 91 out of 102 “other” intervals which have at least 5 CpGs covered by CCS reads in each haplotype are analyzed. The boxes inside the violin plots indicate the 50th percentile (middle line), 25th, and 75th percentile (box). **b** Screenshot of Integrative Genomics Viewer (chr20:60,671,001-60,673,750) on a DMR of SD0651_P1 near the maternally imprinted gene *GNAS*. Red and blue dots represent CpGs with high and low methylation probabilities, respectively.

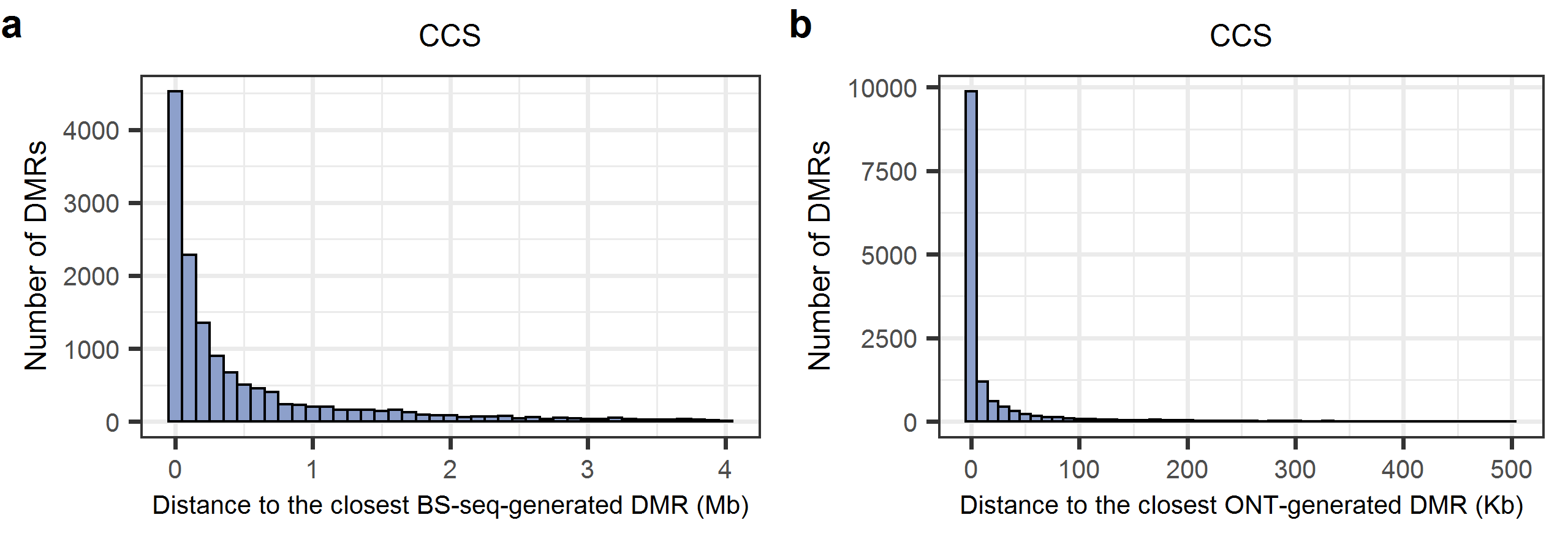

**Supplementary Fig. 9** Distribution of the number of CCS-generated DMRs in terms of distance to the closest BS-seq-generated (**a**) and ONT-generated DMR (**b**) in HG002. ONT: nanopore sequencing.

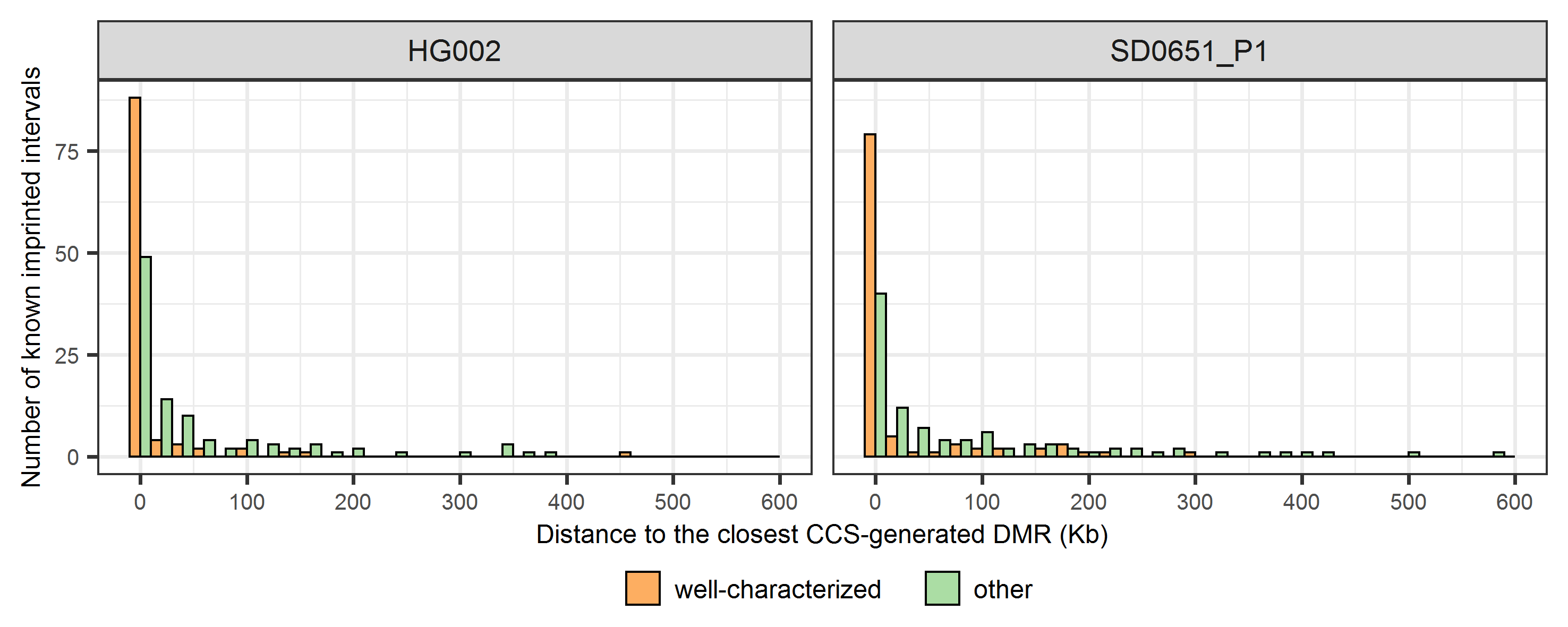

**Supplementary Fig. 10** Distribution of the number of known imprinted intervals in terms of distance to the closest CCS-generated DMR in HG002 and SD0651_P1.

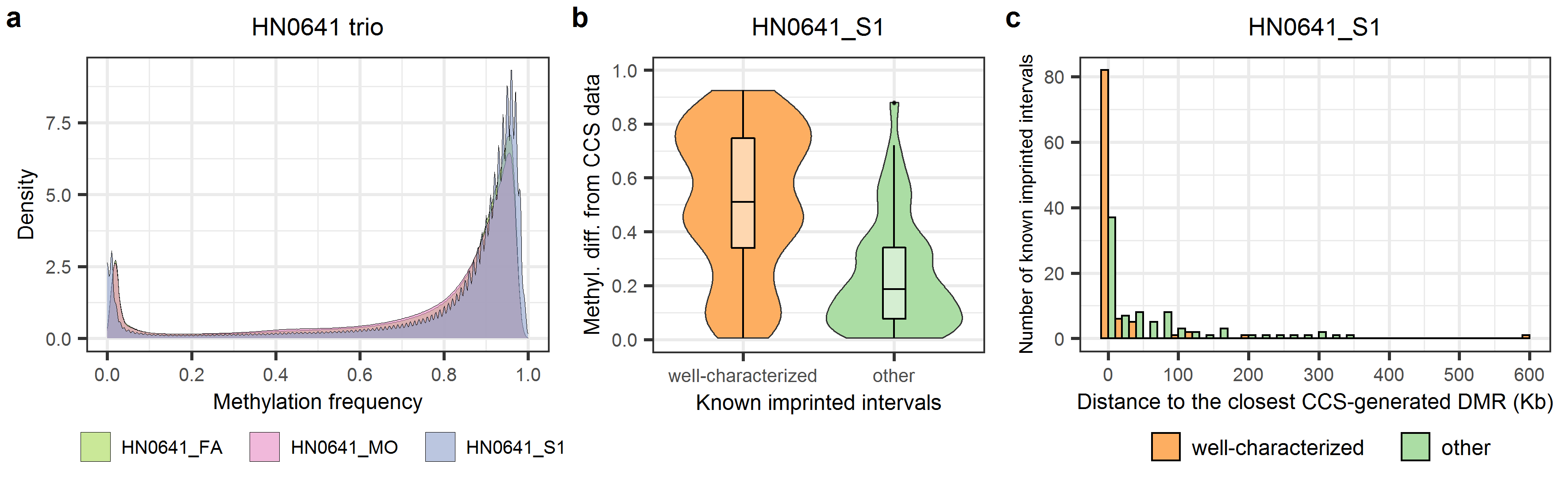

**Supplementary Fig. 11** 5mCpG detection and methylation phasing of the HN0641 family trio using ccsmethphase. **a** Distribution of methylation frequencies of CpGs in HN0641_FA (father), HN0641_MO (mother), and HN0641_S1 (son). **b** Distribution of methylation differences of known imprinted intervals between two haplotypes of HN0641_S1. 93 out of 102 “well-characterized” intervals, and 93 out of 102 “other” intervals which have at least 5 CpGs covered by CCS reads in each haplotype are analyzed. The boxes inside the violin plots indicate the 50th percentile (middle line), 25th, and 75th percentile (box). Dots indicate outliers. **c** Distribution of the number of known imprinted intervals in terms of distance to the closest CCS-generated DMR in HN0641_S1.

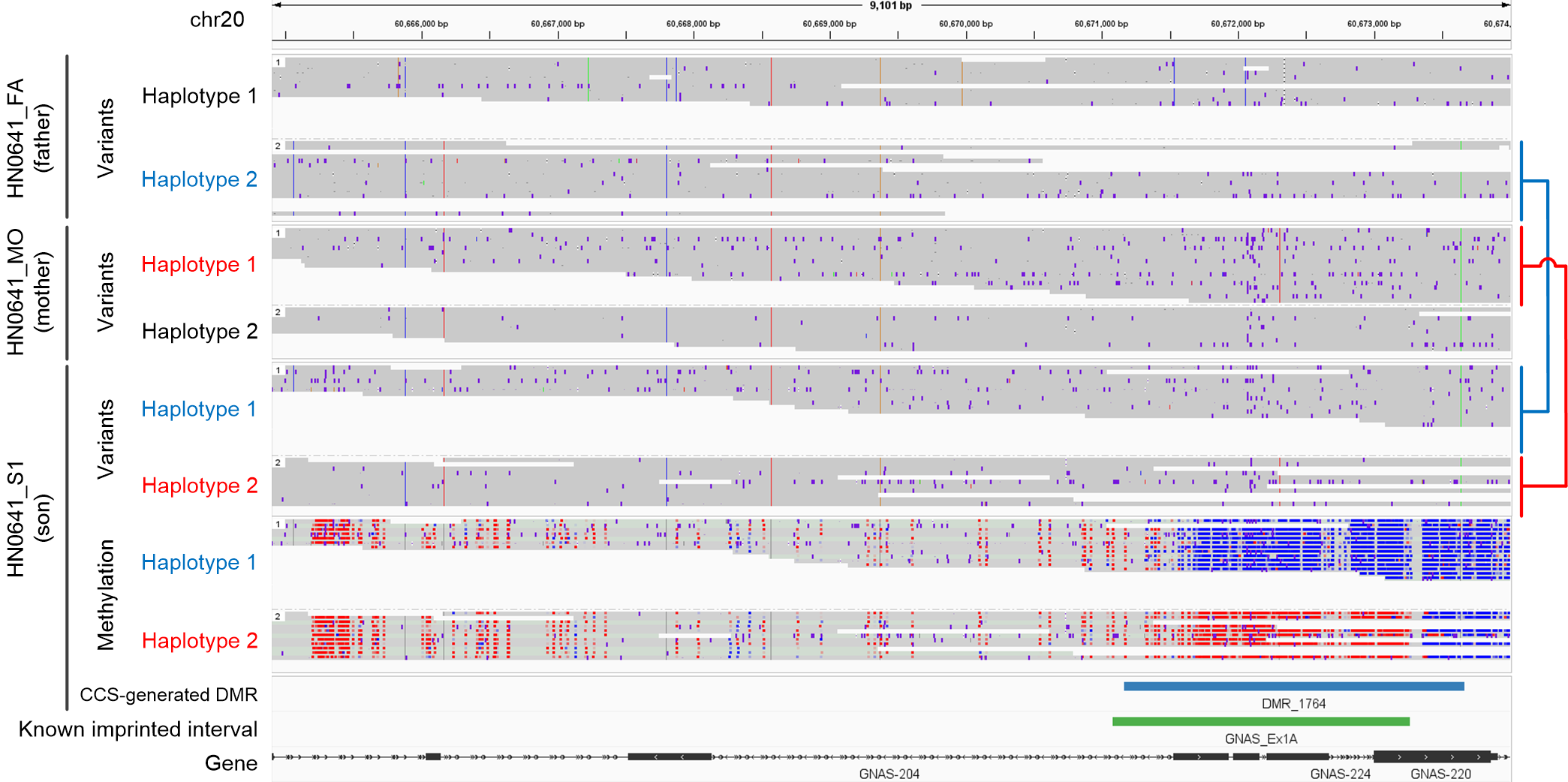

**Supplementary Fig. 12** Screenshot of Integrative Genomics Viewer^1^ (chr20:60,664,900-60,673,999) on a DMR of HN0641_S1 near the **maternally** imprinted gene *GNAS*, showing the variants information of the HN0641 family trio, and the phased methylation information of HN0641_S1. Red and blue dots in the “Methylation” area represent CpGs with high and low methylation probabilities, respectively.

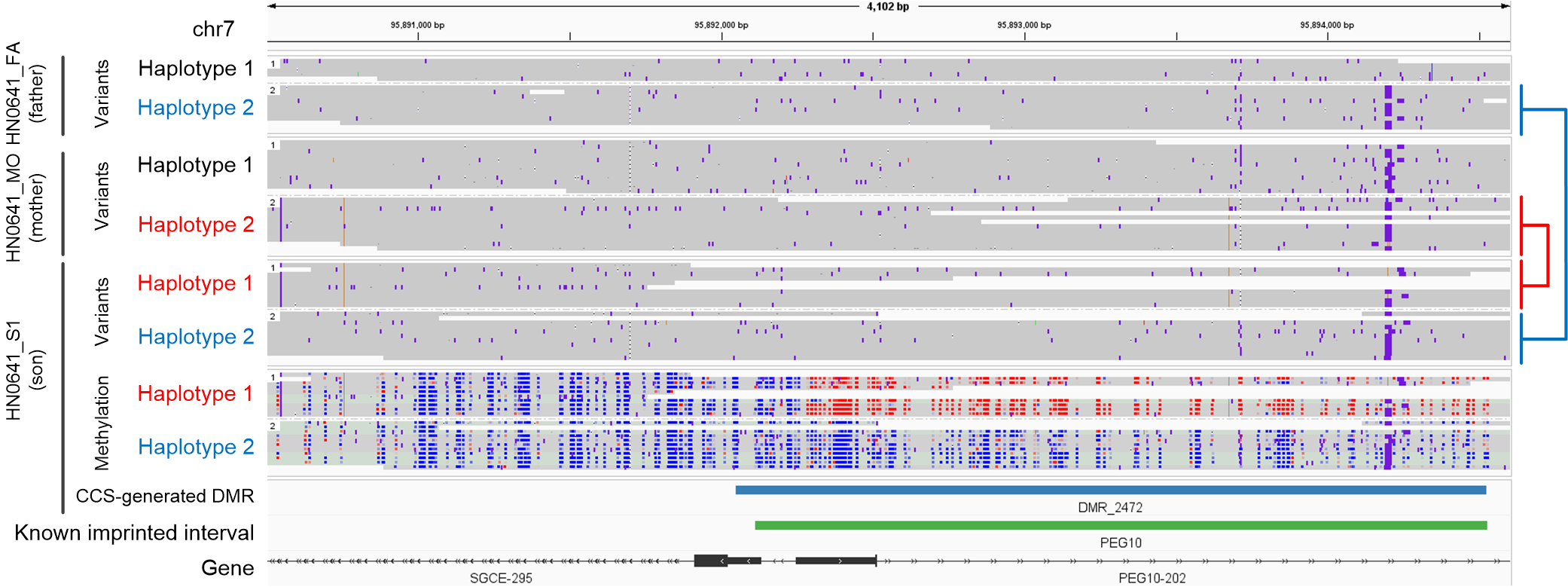

**Supplementary Fig. 13** Screenshot of Integrative Genomics Viewer (chr7:95,890,500-95,894,600) on a DMR of HN0641_S1 near the **maternally** imprinted gene *PEG10*, showing the variants information of the HN0641 family trio, and the phased methylation information of HN0641_S1. Red and blue dots in the “Methylation” area represent CpGs with high and low methylation probabilities, respectively.

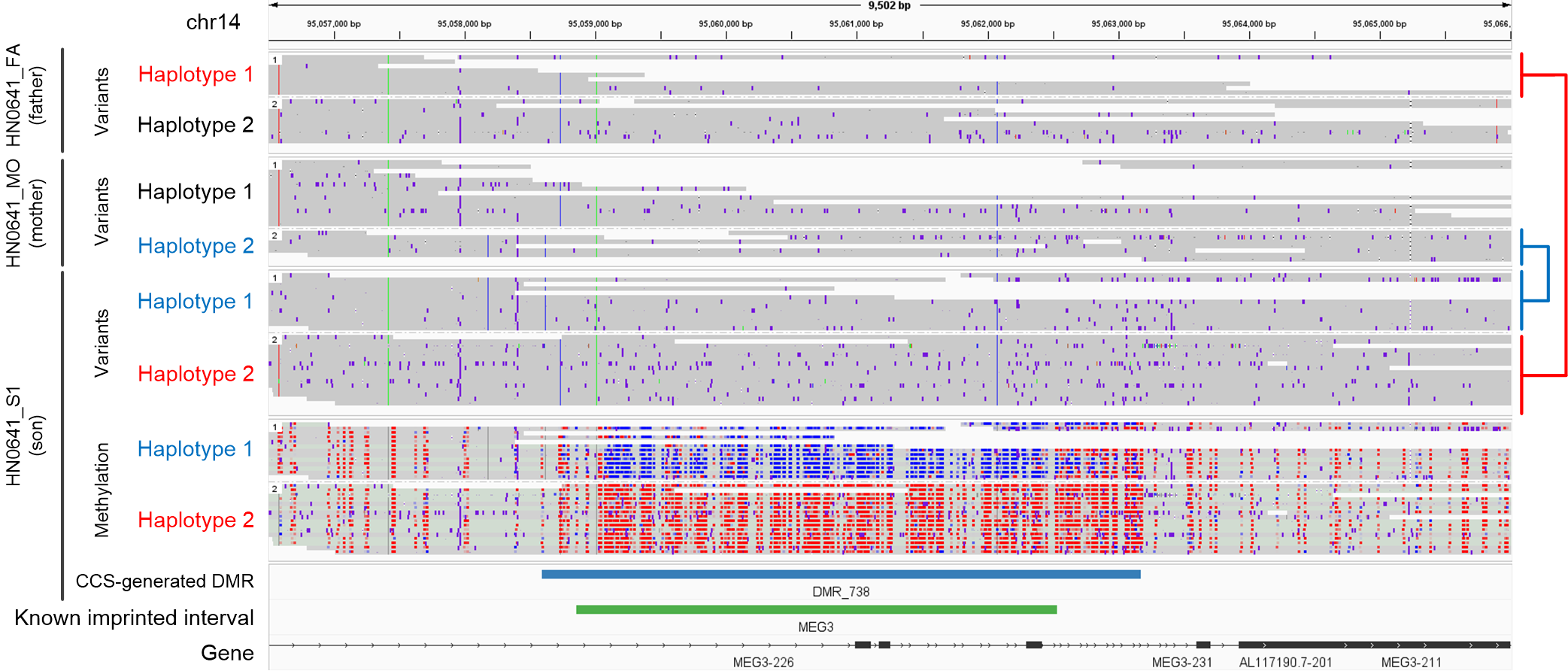

**Supplementary Fig. 14** Screenshot of Integrative Genomics Viewer (chr14:95,056,500-95,066,00) on a DMR of HN0641_S1 near the **paternally** imprinted gene *MEG3*, showing the variants information of the HN0641 family trio, and the phased methylation information of HN0641_S1. Red and blue dots in the “Methylation” area represent CpGs with high and low methylation probabilities, respectively.

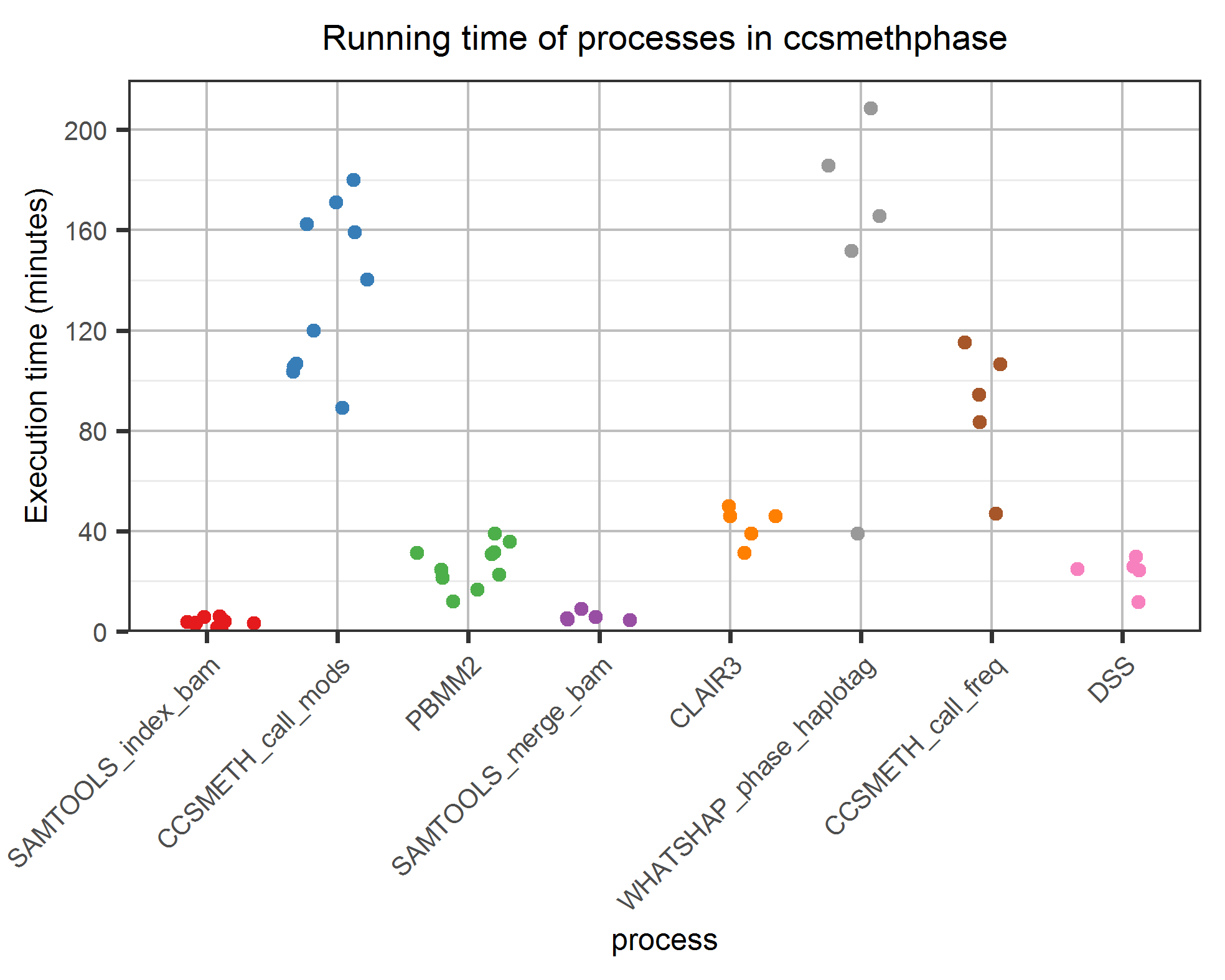

**Supplementary Fig. 15** Running time of 8 main processes in ccsmethphase. 10 SMRT cells of CCS reads (2 SMRT cells for each of the 5 “samples”: HG002 (15Kb), HG002 (20Kb), HG002 (24Kb), CHM13 (20Kb), and SD0651_P1 (15Kb)) were used in this test.

### Supplementary Tables

**Supplementary Table 1** Statistics of PacBio CCS datasets used in this study. read depth: the number of passed subreads for each CCS read; HPRC: Human Pangenome Reference Consortium.

| sample | cell ID | DNA material | sequencing kit | insert  size | NO. of  CCS reads | mean  read length | mean  read depth | source |
| --- | --- | --- | --- | --- | --- | --- | --- | --- |
| M01 | m54276_180627_125201 | M.SssI-treated | sequel I kit 3.0 | - | 423,470 | 440.54 | 42.44 | Tse. *et al.*^2^ |
| W01 | m54276_180627_023725 | PCR-treated | sequel I kit 3.0 | - | 444,310 | 468.12 | 41.66 | Tse. *et al.*^2^ |
| M02 | m64042_190713_204343 | M.SssI-treated | sequel II kit 1.0 | - | 1,144,141 | 4,905.04 | 21.86 | Tse. *et al.*^2^ |
| W02 | m64042_190712_093601 | PCR-treated | sequel II kit 1.0 | - | 1,476,534 | 6,299.35 | 19.23 | Tse. *et al.*^2^ |
| M03 | m64095_200324_133820 | M.SssI-treated | sequel II kit 2.0 | - | 96,933 | 506.72 | 52.85 | Tse. *et al.*^2^ |
| W03 | m64095_200321_184826 | PCR-treated | sequel II kit 2.0 | - | 170,347 | 894.64 | 49.41 | Tse. *et al.*^2^ |
| NA12898 | m64173_220705_133926 | PCR/M.SssI-treated | sequel II kit 2.0 | 10Kb | 2,018,018 | 8,670.84 | 14.12 | in house |
| HG002 | m64012_190921_234837 | native | sequel II kit 2.0 | 15Kb | 2,389,655 | 12,871.53 | 11.09 | HPRC^3^ |
|  | m64012_190920_173625 | native | sequel II kit 2.0 | 15Kb | 2,349,370 | 12,867.23 | 11.05 | HPRC^3^ |
|  | m64015_190920_185703 | native | sequel II kit 2.0 | 15Kb | 2,266,781 | 12,875.45 | 10.97 | HPRC^3^ |
|  | m64008_201124_002822 | native | sequel II kit 2.0 | 15Kb | 2,689,222 | 15,084.49 | 11.29 | Baid *et al.*^4^ |
|  | m64194_201120_222723 | native | sequel II kit 2.0 | 15Kb | 2,606,147 | 15,213.73 | 11.05 | Baid *et al.*^4^ |
|  | m64011_190830_220126 | native | sequel II kit 2.0 | 20Kb | 1,472,376 | 18,521.25 | 9.04 | HPRC^3^ |
|  | m64011_190901_095311 | native | sequel II kit 2.0 | 20Kb | 1,395,877 | 18,516.36 | 9.00 | HPRC^3^ |
|  | m64014_200920_132517 | native | sequel II kit 2.0 | 24Kb | 1,919,428 | 24,160.38 | 7.84 | Baid *et al.*^4^ |
|  | m64179e_200919_061936 | native | sequel II kit 2.0 | 24Kb | 1,742,401 | 24,437.28 | 7.60 | Baid *et al.*^4^ |
| SD0651_P1 | m64114_211125_095059 | native | sequel II kit 2.0 | 15Kb | 1,641,522 | 15,967.21 | 10.37 | in house |
|  | m64242e_211129_171024 | native | sequel II kit 2.0 | 15Kb | 2,178,549 | 16,211.05 | 11.07 | in house |
| CHM13 | m64062_190803_042216 | native | sequel II kit 2.0 | 20Kb | 1,433,166 | 20,760.04 | 7.72 | Nurk *et al.*^5^ |
|  | m64062_190806_063919 | native | sequel II kit 2.0 | 20Kb | 1,045,868 | 20,762.52 | 7.80 | Nurk *et al.*^5^ |
| HN0641_FA | m64242e_211230_172120 | native | sequel II kit 2.0 | 15Kb | 2,105,955 | 15,010.29 | 10.71 | in house |
|  | m64053_220125_054827 | native | sequel II kit 2.0 | 15Kb | 2,379,680 | 15,359.23 | 12.14 | in house |
| HN0641_MO | m64242e_220101_041819 | native | sequel II kit 2.0 | 15Kb | 2,198,341 | 16,179.39 | 10.60 | in house |
|  | m64053_220126_152530 | native | sequel II kit 2.0 | 15Kb | 2,120,743 | 16,363.45 | 11.16 | in house |
| HN0641_S1 | m64242e_220102_151613 | native | sequel II kit 2.0 | 15Kb | 1,851,159 | 16,035.47 | 10.15 | in house |
|  | m64116_220101_042359 | native | sequel II kit 2.0 | 15Kb | 2,254,452 | 16,107.67 | 10.82 | in house |

**Supplementary Table 2** Partition of PacBio long (≥10Kb) CCS reads to evaluate ccsmeth. training_1_: Datasets used to train the model of ccsmeth for read-level 5mCpG prediction; training_2_: Datasets used to train ccsmeth for site-level 5mCpG prediction.

| partition | sample | cell ID | insert size | chromosomes | evaluation |
| --- | --- | --- | --- | --- | --- |
| training_1_ | NA12898 | m64173_220705_133926 | 10Kb | chr1-22 | - |
|  | HG002 | m64012_190921_234837 | 15Kb | chr1-22, chrX, chrY | - |
| training_2_ | HG002 | m64012_190920_173625 | 15Kb | chr1-22, chrX, chrY | - |
|  |  | m64015_190920_185703 | 15Kb | chr1-22, chrX, chrY | - |
| testing | NA12898 | m64173_220705_133926 | 10Kb | chrX | read-level |
|  | HG002 | m64008_201124_002822 | 15Kb | chr1-22, chrX, chrY | read-level, site-level |
|  |  | m64194_201120_222723 | 15Kb | chr1-22, chrX, chrY | read-level, site-level |
|  |  | m64011_190830_220126 | 20Kb | chr1-22, chrX, chrY | read-level, site-level |
|  |  | m64011_190901_095311 | 20Kb | chr1-22, chrX, chrY | read-level, site-level |
|  |  | m64014_200920_132517 | 24Kb | chr1-22, chrX, chrY | read-level, site-level |
|  |  | m64179e_200919_061936 | 24Kb | chr1-22, chrX, chrY | read-level, site-level |
|  | SD0651_P1 | m64114_211125_095059 | 15Kb | chr1-22, chrX, chrY | read-level, site-level |
|  |  | m64242e_211129_171024 | 15Kb | chr1-22, chrX, chrY | read-level, site-level |
|  | CHM13 | m64062_190803_042216 | 20Kb | chr1-22, chrX | site-level |
|  |  | m64062_190806_063919 | 20Kb | chr1-22, chrX | site-level |
|  | HN0641_FA | m64242e_211230_172120 | 15Kb | chr1-22, chrX, chrY | ASM detection |
|  |  | m64053_220125_054827 | 15Kb | chr1-22, chrX, chrY | ASM detection |
|  | HN0641_MO | m64242e_220101_041819 | 15Kb | chr1-22, chrX, chrY | ASM detection |
|  |  | m64053_220126_152530 | 15Kb | chr1-22, chrX, chrY | ASM detection |
|  | HN0641_S1 | m64242e_220102_151613 | 15Kb | chr1-22, chrX, chrY | ASM detection |
|  |  | m64116_220101_042359 | 15Kb | chr1-22, chrX, chrY | ASM detection |

**Supplementary Table 3** Illumina and nanopore datasets used in this study. ONT: Oxford Nanopore Technologies; GIAB: Genome in a Bottle.

| sample | type | (mean)  read length | mean  genome coverage | source |
| --- | --- | --- | --- | --- |
| HG002 | Illumina BS-seq | 2×150 | 117.5× | ONT^6^ |
|  | ONT R9.4.1 | 21,933 | 65.8× | ONT^6^ |
|  | Illumina WGS | 2×250 | 63.1× | GIAB^7^ |
| HG003 | Illumina WGS | 2×250 | 55.7× | GIAB^7^ |
| HG004 | Illumina WGS | 2×250 | 67.9× | GIAB^7^ |
| SD0651_P1 | Illumina BS-seq | 2×150 | 15.7× | in house |
| CHM13 | ONT R9.4.1 | 19,891 | 41.8× | Nurk *et al.*^5^ |

**Supplementary Table 4** Evaluation of ccsmeth and primrose at genome-wide site level against **BS-seq** on **HG002 (15Kb)** dataset. We compared CpGs covered by at least 5 reads in both CCS and BS-seq datasets. For coverage 5×-25×, we subsampled corresponding coverage reads from the total reads, and repeated the subsampling 5 times. Values in the table for coverage 5×-25× are the average of 5 repeated tests. *r*: Pearson correlation; *r^2^*: the coefficient of determination; *ρ*: Spearman correlation; RMSE: root mean square error.

| coverage | method | | *r* | *r^2^* | *ρ* | RMSE |
| --- | --- | --- | --- | --- | --- | --- |
|  | read-level calling | site-level calling |  |  |  |  |
| 5× | primrose | pb-CpG-tools (count) | 0.8145 | 0.6635 | 0.7454 | 0.2058 |
|  | ccsmeth | ccsmeth (count) | 0.8478 | 0.7187 | 0.7741 | 0.1907 |
|  | primrose | pb-CpG-tools (model) | 0.8449 | 0.7139 | 0.7785 | 0.2100 |
|  | primrose | ccsmeth (model) | 0.8784 | 0.7715 | 0.8314 | 0.1694 |
|  | ccsmeth | ccsmeth (model) | 0.8993 | 0.8088 | 0.8510 | 0.1579 |
| 10× | primrose | pb-CpG-tools (count) | 0.8530 | 0.7277 | 0.7821 | 0.1854 |
|  | ccsmeth | ccsmeth (count) | 0.8798 | 0.7741 | 0.8050 | 0.1702 |
|  | primrose | pb-CpG-tools (model) | 0.8864 | 0.7857 | 0.8192 | 0.1781 |
|  | primrose | ccsmeth (model) | 0.9023 | 0.8142 | 0.8554 | 0.1542 |
|  | ccsmeth | ccsmeth (model) | 0.9198 | 0.8459 | 0.8720 | 0.1433 |
| 15× | primrose | pb-CpG-tools (count) | 0.8793 | 0.7732 | 0.8083 | 0.1710 |
|  | ccsmeth | ccsmeth (count) | 0.9008 | 0.8114 | 0.8270 | 0.1559 |
|  | primrose | pb-CpG-tools (model) | 0.9122 | 0.8321 | 0.8462 | 0.1542 |
|  | primrose | ccsmeth (model) | 0.9176 | 0.8419 | 0.8720 | 0.1429 |
|  | ccsmeth | ccsmeth (model) | 0.9326 | 0.8697 | 0.8866 | 0.1327 |
| 20× | primrose | pb-CpG-tools (count) | 0.8953 | 0.8015 | 0.8249 | 0.1622 |
|  | ccsmeth | ccsmeth (count) | 0.9133 | 0.8341 | 0.8410 | 0.1472 |
|  | primrose | pb-CpG-tools (model) | 0.9265 | 0.8585 | 0.8624 | 0.1394 |
|  | primrose | ccsmeth (model) | 0.9269 | 0.8591 | 0.8827 | 0.1353 |
|  | ccsmeth | ccsmeth (model) | 0.9403 | 0.8841 | 0.8960 | 0.1258 |
| 25× | primrose | pb-CpG-tools (count) | 0.9056 | 0.8202 | 0.8360 | 0.1565 |
|  | ccsmeth | ccsmeth (count) | 0.9213 | 0.8489 | 0.8504 | 0.1416 |
|  | primrose | pb-CpG-tools (model) | 0.9354 | 0.8749 | 0.8739 | 0.1297 |
|  | primrose | ccsmeth (model) | 0.9330 | 0.8705 | 0.8899 | 0.1301 |
|  | ccsmeth | ccsmeth (model) | 0.9453 | 0.8935 | 0.9022 | 0.1210 |
| 25.6×(all) | primrose | pb-CpG-tools (count) | 0.9077 | 0.8240 | 0.8383 | 0.1554 |
|  | ccsmeth | ccsmeth (count) | 0.9229 | 0.8518 | 0.8523 | 0.1404 |
|  | primrose | pb-CpG-tools (model) | 0.9371 | 0.8782 | 0.8764 | 0.1278 |
|  | primrose | ccsmeth (model) | 0.9342 | 0.8728 | 0.8913 | 0.1290 |
|  | ccsmeth | ccsmeth (model) | 0.9463 | 0.8955 | 0.9035 | 0.1201 |

**Supplementary Table 5** Evaluation of ccsmeth and primrose at genome-wide site level against **nanopore sequencing** on **HG002 (15Kb)** dataset. We compared CpGs covered by at least 5 reads in both CCS and nanopore datasets. For coverage 5×-25×, we subsampled corresponding coverage reads from the total reads, and repeated the subsampling 5 times. Values in the table for coverage 5×-25× are the average of 5 repeated tests. *r*: Pearson correlation; *r^2^*: the coefficient of determination; *ρ*: Spearman correlation; RMSE: root mean square error.

| coverage | method | | *r* | *r^2^* | *ρ* | RMSE |
| --- | --- | --- | --- | --- | --- | --- |
|  | read-level calling | site-level calling |  |  |  |  |
| 5× | primrose | pb-CpG-tools (count) | 0.7980 | 0.6369 | 0.7378 | 0.2037 |
|  | ccsmeth | ccsmeth (count) | 0.8388 | 0.7037 | 0.7782 | 0.1870 |
|  | primrose | pb-CpG-tools (model) | 0.8248 | 0.6803 | 0.7662 | 0.2228 |
|  | primrose | ccsmeth (model) | 0.8626 | 0.7441 | 0.8135 | 0.1683 |
|  | ccsmeth | ccsmeth (model) | 0.8881 | 0.7887 | 0.8411 | 0.1554 |
| 10× | primrose | pb-CpG-tools (count) | 0.8331 | 0.6941 | 0.7705 | 0.1845 |
|  | ccsmeth | ccsmeth (count) | 0.8685 | 0.7543 | 0.8065 | 0.1674 |
|  | primrose | pb-CpG-tools (model) | 0.8629 | 0.7446 | 0.7994 | 0.1952 |
|  | primrose | ccsmeth (model) | 0.8835 | 0.7806 | 0.8332 | 0.1561 |
|  | ccsmeth | ccsmeth (model) | 0.9062 | 0.8212 | 0.8583 | 0.1433 |
| 15× | primrose | pb-CpG-tools (count) | 0.8557 | 0.7323 | 0.7928 | 0.1714 |
|  | ccsmeth | ccsmeth (count) | 0.8870 | 0.7867 | 0.8262 | 0.1541 |
|  | primrose | pb-CpG-tools (model) | 0.8847 | 0.7828 | 0.8185 | 0.1761 |
|  | primrose | ccsmeth (model) | 0.8959 | 0.8026 | 0.8469 | 0.1477 |
|  | ccsmeth | ccsmeth (model) | 0.9168 | 0.8405 | 0.8704 | 0.1351 |
| 20× | primrose | pb-CpG-tools (count) | 0.8698 | 0.7565 | 0.8075 | 0.1634 |
|  | ccsmeth | ccsmeth (count) | 0.8984 | 0.8070 | 0.8393 | 0.1459 |
|  | primrose | pb-CpG-tools (model) | 0.8970 | 0.8047 | 0.8300 | 0.1641 |
|  | primrose | ccsmeth (model) | 0.9037 | 0.8166 | 0.8560 | 0.1422 |
|  | ccsmeth | ccsmeth (model) | 0.9233 | 0.8525 | 0.8783 | 0.1298 |
| 25× | primrose | pb-CpG-tools (count) | 0.8794 | 0.7733 | 0.8178 | 0.1581 |
|  | ccsmeth | ccsmeth (count) | 0.9060 | 0.8209 | 0.8485 | 0.1403 |
|  | primrose | pb-CpG-tools (model) | 0.9049 | 0.8189 | 0.8395 | 0.1561 |
|  | primrose | ccsmeth (model) | 0.9091 | 0.8264 | 0.8624 | 0.1382 |
|  | ccsmeth | ccsmeth (model) | 0.9278 | 0.8608 | 0.8839 | 0.1260 |
| 25.6×(all) | primrose | pb-CpG-tools (count) | 0.8813 | 0.7768 | 0.8200 | 0.1569 |
|  | ccsmeth | ccsmeth (count) | 0.9076 | 0.8238 | 0.8504 | 0.1391 |
|  | primrose | pb-CpG-tools (model) | 0.9065 | 0.8218 | 0.8417 | 0.1545 |
|  | primrose | ccsmeth (model) | 0.9102 | 0.8284 | 0.8637 | 0.1373 |
|  | ccsmeth | ccsmeth (model) | 0.9287 | 0.8626 | 0.8850 | 0.1252 |

**Supplementary Table 6** Evaluation of ccsmeth and primrose at genome-wide site level against **BS-seq** on **HG002 (20Kb)** dataset. We compared CpGs covered by at least 5 reads in both CCS and BS-seq datasets. For coverage 5×-15×, we subsampled corresponding coverage reads from the total reads, and repeated the subsampling 5 times. Values in the table for coverage 5×-15× are the average of 5 repeated tests. *r*: Pearson correlation; *r^2^*: the coefficient of determination; *ρ*: Spearman correlation; RMSE: root mean square error.

| coverage | method | | *r* | *r^2^* | *ρ* | RMSE |
| --- | --- | --- | --- | --- | --- | --- |
|  | read-level calling | site-level calling |  |  |  |  |
| 5× | primrose | pb-CpG-tools (count) | 0.7800 | 0.6084 | 0.7060 | 0.2205 |
|  | ccsmeth | ccsmeth (count) | 0.8253 | 0.6811 | 0.7449 | 0.2002 |
|  | primrose | pb-CpG-tools (model) | 0.8165 | 0.6667 | 0.7425 | 0.2266 |
|  | primrose | ccsmeth (model) | 0.8406 | 0.7067 | 0.7806 | 0.1906 |
|  | ccsmeth | ccsmeth (model) | 0.8842 | 0.7817 | 0.8283 | 0.1636 |
| 10× | primrose | pb-CpG-tools (count) | 0.8247 | 0.6802 | 0.7462 | 0.2008 |
|  | ccsmeth | ccsmeth (count) | 0.8606 | 0.7407 | 0.7763 | 0.1805 |
|  | primrose | pb-CpG-tools (model) | 0.8653 | 0.7487 | 0.7864 | 0.1931 |
|  | primrose | ccsmeth (model) | 0.8726 | 0.7614 | 0.8071 | 0.1748 |
|  | ccsmeth | ccsmeth (model) | 0.9083 | 0.8251 | 0.8499 | 0.1482 |
| 15× | primrose | pb-CpG-tools (count) | 0.8549 | 0.7308 | 0.7750 | 0.1866 |
|  | ccsmeth | ccsmeth (count) | 0.8829 | 0.7794 | 0.7989 | 0.1667 |
|  | primrose | pb-CpG-tools (model) | 0.8947 | 0.8004 | 0.8163 | 0.1686 |
|  | primrose | ccsmeth (model) | 0.8924 | 0.7964 | 0.8263 | 0.1629 |
|  | ccsmeth | ccsmeth (model) | 0.9224 | 0.8509 | 0.8653 | 0.1372 |
| 17.0×(all) | primrose | pb-CpG-tools (count) | 0.8648 | 0.7479 | 0.7849 | 0.1819 |
|  | ccsmeth | ccsmeth (count) | 0.8900 | 0.7922 | 0.8066 | 0.1622 |
|  | primrose | pb-CpG-tools (model) | 0.9038 | 0.8168 | 0.8265 | 0.1603 |
|  | primrose | ccsmeth (model) | 0.8991 | 0.8084 | 0.8334 | 0.1586 |
|  | ccsmeth | ccsmeth (model) | 0.9271 | 0.8594 | 0.8707 | 0.1333 |

**Supplementary Table 7** Evaluation of ccsmeth and primrose at genome-wide site level against **nanopore sequencing** on **HG002 (20Kb)** dataset. We compared CpGs covered by at least 5 reads in both CCS and nanopore datasets. For coverage 5×-15×, we subsampled corresponding coverage reads from the total reads, and repeated the subsampling 5 times. Values in the table for coverage 5×-15× are the average of 5 repeated tests. *r*: Pearson correlation; *r^2^*: the coefficient of determination; *ρ*: Spearman correlation; RMSE: root mean square error.

| coverage | method | | *r* | *r^2^* | *ρ* | RMSE |
| --- | --- | --- | --- | --- | --- | --- |
|  | read-level calling | site-level calling |  |  |  |  |
| 5× | primrose | pb-CpG-tools (count) | 0.7647 | 0.5847 | 0.7019 | 0.2159 |
|  | ccsmeth | ccsmeth (count) | 0.8192 | 0.6711 | 0.7554 | 0.1950 |
|  | primrose | pb-CpG-tools (model) | 0.7967 | 0.6348 | 0.7330 | 0.2376 |
|  | primrose | ccsmeth (model) | 0.8262 | 0.6827 | 0.7691 | 0.1861 |
|  | ccsmeth | ccsmeth (model) | 0.8749 | 0.7655 | 0.8242 | 0.1608 |
| 10× | primrose | pb-CpG-tools (count) | 0.8061 | 0.6498 | 0.7389 | 0.1968 |
|  | ccsmeth | ccsmeth (count) | 0.8522 | 0.7263 | 0.7851 | 0.1759 |
|  | primrose | pb-CpG-tools (model) | 0.8422 | 0.7092 | 0.7700 | 0.2078 |
|  | primrose | ccsmeth (model) | 0.8549 | 0.7308 | 0.792 | 0.1725 |
|  | ccsmeth | ccsmeth (model) | 0.8967 | 0.8040 | 0.8424 | 0.1481 |
| 15× | primrose | pb-CpG-tools (count) | 0.8327 | 0.6934 | 0.7645 | 0.1834 |
|  | ccsmeth | ccsmeth (count) | 0.8723 | 0.7608 | 0.8061 | 0.1628 |
|  | primrose | pb-CpG-tools (model) | 0.8679 | 0.7532 | 0.7929 | 0.1873 |
|  | primrose | ccsmeth (model) | 0.8718 | 0.7600 | 0.8087 | 0.1627 |
|  | ccsmeth | ccsmeth (model) | 0.9087 | 0.8258 | 0.8555 | 0.1395 |
| 17.0×(all) | primrose | pb-CpG-tools (count) | 0.8416 | 0.7084 | 0.7735 | 0.1789 |
|  | ccsmeth | ccsmeth (count) | 0.8788 | 0.7724 | 0.8135 | 0.1584 |
|  | primrose | pb-CpG-tools (model) | 0.8758 | 0.7671 | 0.8008 | 0.1804 |
|  | primrose | ccsmeth (model) | 0.8776 | 0.7701 | 0.8152 | 0.1591 |
|  | ccsmeth | ccsmeth (model) | 0.9127 | 0.8330 | 0.8602 | 0.1365 |

**Supplementary Table 8** Evaluation of ccsmeth and primrose at genome-wide site level against **BS-seq** on **HG002 (24Kb)** dataset. We compared CpGs covered by at least 5 reads in both CCS and BS-seq datasets. For coverage 5×-25×, we subsampled corresponding coverage reads from the total reads, and repeated the subsampling 5 times. Values in the table for coverage 5×-25× are the average of 5 repeated tests. *r*: Pearson correlation; *r^2^*: the coefficient of determination; *ρ*: Spearman correlation; RMSE: root mean square error.

| coverage | method | | *r* | *r^2^* | *ρ* | RMSE |
| --- | --- | --- | --- | --- | --- | --- |
|  | read-level calling | site-level calling |  |  |  |  |
| 5× | primrose | pb-CpG-tools (count) | 0.7818 | 0.6112 | 0.7156 | 0.2217 |
|  | ccsmeth | ccsmeth (count) | 0.8249 | 0.6805 | 0.7524 | 0.2026 |
|  | primrose | pb-CpG-tools (model) | 0.8247 | 0.6801 | 0.7571 | 0.2221 |
|  | primrose | ccsmeth (model) | 0.8586 | 0.7372 | 0.8097 | 0.1810 |
|  | ccsmeth | ccsmeth (model) | 0.8864 | 0.7857 | 0.8360 | 0.1645 |
| 10× | primrose | pb-CpG-tools (count) | 0.8246 | 0.6799 | 0.7570 | 0.2017 |
|  | ccsmeth | ccsmeth (count) | 0.8605 | 0.7404 | 0.7873 | 0.1822 |
|  | primrose | pb-CpG-tools (model) | 0.8696 | 0.7562 | 0.8006 | 0.1898 |
|  | primrose | ccsmeth (model) | 0.8846 | 0.7826 | 0.8363 | 0.1660 |
|  | ccsmeth | ccsmeth (model) | 0.9087 | 0.8257 | 0.8591 | 0.1494 |
| 15× | primrose | pb-CpG-tools (count) | 0.8545 | 0.7301 | 0.7862 | 0.1877 |
|  | ccsmeth | ccsmeth (count) | 0.8841 | 0.7817 | 0.8118 | 0.1682 |
|  | primrose | pb-CpG-tools (model) | 0.8983 | 0.8070 | 0.8298 | 0.1652 |
|  | primrose | ccsmeth (model) | 0.9017 | 0.8131 | 0.8546 | 0.1546 |
|  | ccsmeth | ccsmeth (model) | 0.9230 | 0.8519 | 0.8749 | 0.1381 |
| 20× | primrose | pb-CpG-tools (count) | 0.8729 | 0.7620 | 0.8047 | 0.1793 |
|  | ccsmeth | ccsmeth (count) | 0.8985 | 0.8073 | 0.8273 | 0.1597 |
|  | primrose | pb-CpG-tools (model) | 0.9148 | 0.8368 | 0.8479 | 0.1496 |
|  | primrose | ccsmeth (model) | 0.9124 | 0.8324 | 0.8664 | 0.1468 |
|  | ccsmeth | ccsmeth (model) | 0.9318 | 0.8682 | 0.8850 | 0.1305 |
| 25× | primrose | pb-CpG-tools (count) | 0.8851 | 0.7834 | 0.8170 | 0.1738 |
|  | ccsmeth | ccsmeth (count) | 0.9078 | 0.8241 | 0.8377 | 0.1541 |
|  | primrose | pb-CpG-tools (model) | 0.9250 | 0.8556 | 0.8608 | 0.1394 |
|  | primrose | ccsmeth (model) | 0.9194 | 0.8453 | 0.8745 | 0.1413 |
|  | ccsmeth | ccsmeth (model) | 0.9376 | 0.8790 | 0.8918 | 0.1252 |
| 28.4×(all) | primrose | pb-CpG-tools (count) | 0.8923 | 0.7962 | 0.8244 | 0.1705 |
|  | ccsmeth | ccsmeth (count) | 0.9132 | 0.8340 | 0.8439 | 0.1508 |
|  | primrose | pb-CpG-tools (model) | 0.9309 | 0.8666 | 0.8692 | 0.1333 |
|  | primrose | ccsmeth (model) | 0.9237 | 0.8532 | 0.8794 | 0.1378 |
|  | ccsmeth | ccsmeth (model) | 0.9410 | 0.8855 | 0.8960 | 0.1220 |

**Supplementary Table 9** Evaluation of ccsmeth and primrose at genome-wide site level against **nanopore sequencing** on **HG002 (24Kb)** dataset. We compared CpGs covered by at least 5 reads in both CCS and nanopore datasets. For coverage 5×-25×, we subsampled corresponding coverage reads from the total reads, and repeated the subsampling 5 times. Values in the table for coverage 5×-25× are the average of 5 repeated tests. *r*: Pearson correlation; *r^2^*: the coefficient of determination; *ρ*: Spearman correlation; RMSE: root mean square error.

| coverage | method | | *r* | *r^2^* | *ρ* | RMSE |
| --- | --- | --- | --- | --- | --- | --- |
|  | read-level calling | site-level calling |  |  |  |  |
| 5× | primrose | pb-CpG-tools (count) | 0.7660 | 0.5868 | 0.7090 | 0.2163 |
|  | ccsmeth | ccsmeth (count) | 0.8165 | 0.6667 | 0.7575 | 0.1964 |
|  | primrose | pb-CpG-tools (model) | 0.8042 | 0.6467 | 0.7456 | 0.2341 |
|  | primrose | ccsmeth (model) | 0.8432 | 0.7110 | 0.7932 | 0.1782 |
|  | ccsmeth | ccsmeth (model) | 0.8752 | 0.7659 | 0.8267 | 0.1616 |
| 10× | primrose | pb-CpG-tools (count) | 0.8058 | 0.6493 | 0.7472 | 0.1969 |
|  | ccsmeth | ccsmeth (count) | 0.8501 | 0.7226 | 0.7904 | 0.1765 |
|  | primrose | pb-CpG-tools (model) | 0.8462 | 0.7161 | 0.7823 | 0.2054 |
|  | primrose | ccsmeth (model) | 0.8663 | 0.7504 | 0.8157 | 0.1659 |
|  | ccsmeth | ccsmeth (model) | 0.8952 | 0.8014 | 0.8461 | 0.1491 |
| 15× | primrose | pb-CpG-tools (count) | 0.8324 | 0.6928 | 0.7731 | 0.1836 |
|  | ccsmeth | ccsmeth (count) | 0.8717 | 0.7598 | 0.8132 | 0.1628 |
|  | primrose | pb-CpG-tools (model) | 0.8715 | 0.7595 | 0.8043 | 0.1849 |
|  | primrose | ccsmeth (model) | 0.8807 | 0.7756 | 0.8313 | 0.1571 |
|  | ccsmeth | ccsmeth (model) | 0.9076 | 0.8237 | 0.8596 | 0.1403 |
| 20× | primrose | pb-CpG-tools (count) | 0.8492 | 0.7211 | 0.7900 | 0.1754 |
|  | ccsmeth | ccsmeth (count) | 0.8852 | 0.7836 | 0.8280 | 0.1543 |
|  | primrose | pb-CpG-tools (model) | 0.8862 | 0.7853 | 0.8183 | 0.1718 |
|  | primrose | ccsmeth (model) | 0.8900 | 0.7920 | 0.8417 | 0.1511 |
|  | ccsmeth | ccsmeth (model) | 0.9154 | 0.8380 | 0.8684 | 0.1343 |
| 25× | primrose | pb-CpG-tools (count) | 0.8605 | 0.7405 | 0.8015 | 0.1700 |
|  | ccsmeth | ccsmeth (count) | 0.8942 | 0.7996 | 0.8382 | 0.1486 |
|  | primrose | pb-CpG-tools (model) | 0.8955 | 0.8020 | 0.8293 | 0.1631 |
|  | primrose | ccsmeth (model) | 0.8963 | 0.8033 | 0.8488 | 0.1469 |
|  | ccsmeth | ccsmeth (model) | 0.9207 | 0.8477 | 0.8744 | 0.1301 |
| 28.4×(all) | primrose | pb-CpG-tools (count) | 0.8674 | 0.7524 | 0.8086 | 0.1668 |
|  | ccsmeth | ccsmeth (count) | 0.8996 | 0.8093 | 0.8445 | 0.1452 |
|  | primrose | pb-CpG-tools (model) | 0.9011 | 0.8119 | 0.8370 | 0.1579 |
|  | primrose | ccsmeth (model) | 0.9002 | 0.8103 | 0.8532 | 0.1442 |
|  | ccsmeth | ccsmeth (model) | 0.9240 | 0.8537 | 0.8781 | 0.1275 |

**Supplementary Table 10** Evaluation of ccsmeth and primrose at genome-wide site level against **BS-seq** on **SD0651_P1 (15Kb)** dataset. We compared CpGs covered by at least 5 reads in both CCS and BS-seq datasets. For coverage 5×-15×, we subsampled corresponding coverage reads from the total reads, and repeated the subsampling 5 times. Values in the table for coverage 5×-15× are the average of 5 repeated tests. *r*: Pearson correlation; *r^2^*: the coefficient of determination; *ρ*: Spearman correlation; RMSE: root mean square error.

| coverage | method | | *r* | *r^2^* | *ρ* | RMSE |
| --- | --- | --- | --- | --- | --- | --- |
|  | read-level calling | site-level calling |  |  |  |  |
| 5× | primrose | pb-CpG-tools (count) | 0.6670 | 0.4449 | 0.3993 | 0.2279 |
|  | ccsmeth | ccsmeth (count) | 0.7160 | 0.5127 | 0.4463 | 0.2095 |
|  | primrose | pb-CpG-tools (model) | 0.7738 | 0.5988 | 0.4598 | 0.1772 |
|  | primrose | ccsmeth (model) | 0.7528 | 0.5667 | 0.4372 | 0.1897 |
|  | ccsmeth | ccsmeth (model) | 0.8233 | 0.6778 | 0.5012 | 0.1554 |
| 10× | primrose | pb-CpG-tools (count) | 0.7218 | 0.5210 | 0.4305 | 0.2095 |
|  | ccsmeth | ccsmeth (count) | 0.7633 | 0.5826 | 0.4742 | 0.1913 |
|  | primrose | pb-CpG-tools (model) | 0.8213 | 0.6745 | 0.4807 | 0.1568 |
|  | primrose | ccsmeth (model) | 0.7873 | 0.6198 | 0.4551 | 0.1781 |
|  | ccsmeth | ccsmeth (model) | 0.8506 | 0.7235 | 0.5210 | 0.1437 |
| 15× | primrose | pb-CpG-tools (count) | 0.7598 | 0.5773 | 0.4542 | 0.1969 |
|  | ccsmeth | ccsmeth (count) | 0.7940 | 0.6304 | 0.4964 | 0.1792 |
|  | primrose | pb-CpG-tools (model) | 0.8501 | 0.7226 | 0.4930 | 0.1416 |
|  | primrose | ccsmeth (model) | 0.8090 | 0.6544 | 0.4688 | 0.1691 |
|  | ccsmeth | ccsmeth (model) | 0.8653 | 0.7488 | 0.5358 | 0.1359 |
| 19.6×(all) | primrose | pb-CpG-tools (count) | 0.7839 | 0.6146 | 0.4707 | 0.1893 |
|  | ccsmeth | ccsmeth (count) | 0.8128 | 0.6607 | 0.5117 | 0.1720 |
|  | primrose | pb-CpG-tools (model) | 0.8702 | 0.7572 | 0.5093 | 0.1274 |
|  | primrose | ccsmeth (model) | 0.8239 | 0.6788 | 0.4800 | 0.1624 |
|  | ccsmeth | ccsmeth (model) | 0.8750 | 0.7656 | 0.5461 | 0.1303 |

**Supplementary Table 11** Evaluation of ccsmeth and primrose at genome-wide site level against **nanopore sequencing** on **CHM13 (20Kb)** dataset. We compared CpGs covered by at least 5 reads in both CCS and nanopore datasets. For coverage 5×-15×, we subsampled corresponding coverage reads from the total reads, and repeated the subsampling 5 times. Values in the table for coverage 5×-15× are the average of 5 repeated tests. *r*: Pearson correlation; *r^2^*: the coefficient of determination; *ρ*: Spearman correlation; RMSE: root mean square error.

| coverage | method | | *r* | *r^2^* | *ρ* | RMSE |
| --- | --- | --- | --- | --- | --- | --- |
|  | read-level calling | site-level calling |  |  |  |  |
| 5× | primrose | pb-CpG-tools (count) | 0.7941 | 0.6305 | 0.7621 | 0.2192 |
|  | ccsmeth | ccsmeth (count) | 0.8359 | 0.6987 | 0.8124 | 0.2000 |
|  | primrose | pb-CpG-tools (model) | 0.8332 | 0.6943 | 0.8029 | 0.2334 |
|  | primrose | ccsmeth (model) | 0.8698 | 0.7566 | 0.8248 | 0.1759 |
|  | ccsmeth | ccsmeth (model) | 0.8989 | 0.8080 | 0.8643 | 0.1587 |
| 10× | primrose | pb-CpG-tools (count) | 0.8357 | 0.6984 | 0.8032 | 0.1967 |
|  | ccsmeth | ccsmeth (count) | 0.8692 | 0.7556 | 0.8456 | 0.1779 |
|  | primrose | pb-CpG-tools (model) | 0.8787 | 0.7721 | 0.8427 | 0.1966 |
|  | primrose | ccsmeth (model) | 0.8939 | 0.7991 | 0.8479 | 0.1608 |
|  | ccsmeth | ccsmeth (model) | 0.9181 | 0.8430 | 0.8848 | 0.1448 |
| 15× | primrose | pb-CpG-tools (count) | 0.8639 | 0.7462 | 0.8286 | 0.1810 |
|  | ccsmeth | ccsmeth (count) | 0.8911 | 0.7940 | 0.8667 | 0.1622 |
|  | primrose | pb-CpG-tools (model) | 0.9050 | 0.8191 | 0.8640 | 0.1714 |
|  | primrose | ccsmeth (model) | 0.9087 | 0.8257 | 0.8605 | 0.1499 |
|  | ccsmeth | ccsmeth (model) | 0.9299 | 0.8647 | 0.8962 | 0.1350 |
| 16.5×(all) | primrose | pb-CpG-tools (count) | 0.8711 | 0.7589 | 0.8350 | 0.1770 |
|  | ccsmeth | ccsmeth (count) | 0.8966 | 0.8039 | 0.8720 | 0.1582 |
|  | primrose | pb-CpG-tools (model) | 0.9112 | 0.8303 | 0.8688 | 0.1650 |
|  | primrose | ccsmeth (model) | 0.9124 | 0.8325 | 0.8636 | 0.1469 |
|  | ccsmeth | ccsmeth (model) | 0.9328 | 0.8701 | 0.8990 | 0.1324 |

**Supplementary Table 12** Comparing CCS with BS-seq and nanopore sequencing on predicting site-level methylation frequencies of HG002 haplotypes phased by Illumina trio data. CpGs in haplotypes of autosomes were used for evaluation. *r*: Pearson correlation; *r^2^*: the coefficient of determination; *ρ*: Spearman correlation; RMSE: root mean square error; ONT: nanopore sequencing.

| benchmark | haplotype | *r* | *r^2^* | *ρ* | RMSE |
| --- | --- | --- | --- | --- | --- |
| BS-seq | maternal | 0.9321 | 0.8689 | 0.8505 | 0.1366 |
|  | paternal | 0.9322 | 0.8689 | 0.8507 | 0.1365 |
| ONT | maternal | 0.9403 | 0.8842 | 0.8623 | 0.1215 |
|  | paternal | 0.9404 | 0.8844 | 0.8625 | 0.1214 |

**Supplementary Table 13** Comparing CCS (ccsmeth) with BS-seq (Bismark) and nanopore sequencing (DeepSignal2) on predicting site-level methylation frequencies in repetitive genomic regions using HG002 data. *r*: Pearson correlation; *r^2^*: the coefficient of determination; *ρ*: Spearman correlation; RMSE: root mean square error; ONT: nanopore sequencing; RepeatMasker: repetitive genomic elements annotated by RepeatMasker; SDs: segmental duplications; cenSats: peri/centromeric satellites.

| region | benchmark | *r* | *r^2^* | *ρ* | RMSE |
| --- | --- | --- | --- | --- | --- |
| RepeatMasker | BS-seq | 0.9540 | 0.9101 | 0.9102 | 0.1055 |
|  | ONT | 0.9358 | 0.8758 | 0.8902 | 0.1138 |
| SDs | BS-seq | 0.9208 | 0.8479 | 0.8770 | 0.1370 |
|  | ONT | 0.9087 | 0.8257 | 0.8791 | 0.1308 |
| cenSats | BS-seq | 0.8822 | 0.7783 | 0.8462 | 0.1584 |
|  | ONT | 0.8572 | 0.7349 | 0.8327 | 0.1606 |

**Supplementary Table 14** Comparing CCS (ccsmeth) with BS-seq (Bismark) and nanopore sequencing (DeepSignal2) on predicting site-level methylation frequencies in repetitive genomic regions of HG002 haplotypes phased by Illumina trio data. *r*: Pearson correlation; *r^2^*: the coefficient of determination; *ρ*: Spearman correlation; RMSE: root mean square error; ONT: nanopore sequencing; RepeatMasker: repetitive genomic elements annotated by RepeatMasker; SDs: segmental duplications; cenSats: peri/centromeric satellites.

| region | benchmark | haplotype | *r* | *r^2^* | *ρ* | RMSE |
| --- | --- | --- | --- | --- | --- | --- |
| RepeatMasker | BS-seq | maternal | 0.9283 | 0.8617 | 0.8429 | 0.1370 |
|  |  | paternal | 0.9285 | 0.8621 | 0.8432 | 0.1369 |
|  | ONT | maternal | 0.9365 | 0.8771 | 0.8546 | 0.1221 |
|  |  | paternal | 0.9368 | 0.8775 | 0.8550 | 0.1218 |
| SDs | BS-seq | maternal | 0.9053 | 0.8196 | 0.8332 | 0.1558 |
|  |  | paternal | 0.9007 | 0.8113 | 0.8311 | 0.1599 |
|  | ONT | maternal | 0.9175 | 0.8419 | 0.8490 | 0.1377 |
|  |  | paternal | 0.9160 | 0.8390 | 0.8520 | 0.1389 |
| cenSats | BS-seq | maternal | 0.8907 | 0.7933 | 0.8340 | 0.1633 |
|  |  | paternal | 0.8925 | 0.7966 | 0.8378 | 0.1628 |
|  | ONT | maternal | 0.9023 | 0.8141 | 0.8553 | 0.1489 |
|  |  | paternal | 0.9066 | 0.8219 | 0.8612 | 0.1458 |

**Supplementary Table 15** Computing resources used for evaluating the running time of the processes in ccsmethphase.

| process | server | No. of CPU cores | No. of GPU cards |
| --- | --- | --- | --- |
| *SAMTOOLS_index_bam* | Server-CPU | 40 | - |
| *CCSMETH_call_mods* | Server-GPU | 40 | 2 |
| *PBMM2* | Server-CPU | 40 | - |
| *SAMTOOLS_merge_bam* | Server-CPU | 40 | - |
| *CLAIR3* | Server-CPU | 40 | - |
| *WHATSHAP_phase_haplotag* | Server-CPU | 10 | - |
| *CCSMETH_call_freq* | Server-CPU | 40 | - |
| *DSS* | Server-CPU | 40 | - |

### Supplementary Notes

**Supplementary Note 1** **Pipeline for haplotype-aware methylation calling using Illumina whole-genome sequencing (WGS) trio data and BS-seq data**

To evaluate the methylation phasing pipeline on CCS data, we performed haplotype-aware methylation calling using WGS and BS-seq reads of HG002 as the benchmark (Supplementary Fig. 2). In this pipeline, we used SNPsplit (version 0.5.0) ^8^ to assign BS-seq reads to the haplotypes of HG002. SNPsplit requires information of heterozygous SNVs of HG002 and the origin of these SNVs for accurate reads alignment and splitting. Thus, we downloaded the Illumina WGS reads of AshkenazimTrio: HG003 is the father, HG004 is the mother, HG002 is the son. We used BWA-MEM (version 0.7.17-r1194-dirty)^9^ to align the WGS reads, and then used DeepTrio^10^ (version 1.3.0) to call SNVs for HG002, HG003, and HG004. Then, we used the following rules as in NanoMethPhase^11^ to phase heterozygous SNVs in autosomes of HG002:

$\left\{ \begin{aligned} &\mathrm{Haplotype}\left( S \right)=1, if S\in maternal SNVs and S\notin\mathrm{paternal}\mathrm{SNVs} and S\in child's heterozygous SNVs \\ &Haplotype\left( S \right)=1, else if S\in maternal homozygous SNVs and S\notin\mathrm{paternal} homozygous SNVs and S\in child's heterozygous SNVs \\ &Haplotype\left( S \right)=2, else if S\notin maternal SNVs and S\in\mathrm{paternal}\mathrm{SNVs} and S\in child's heterozygous SNVs \\ &Haplotype\left( S \right)=2, else if S\notin maternal homozygous SNVs and S\in\mathrm{paternal} homozygous SNVs and S\in child's heterozygous SNVs \end{aligned} \right.$ (**1**)

where *S* represents an SNV. After phasing, we generated a chromosome-level SNV phasing result (*i.e.*, all heterozygous SNVs of HG002 inherited from HG004 (mother) were assigned to Haplotype 1, and all heterozygous SNVs inherited from HG003 (father) were assigned to Haplotype 2).

We used Bismark^12^ to align BS-seq reads to genome reference. Then, we used SNPsplit to assign the aligned BS-seq reads to haplotypes. We got the methylation profile of each haplotype using Bismark. At last, we got differentially methylated regions (DMRs) of the two haplotypes using DSS (version 2.44.0)^13^ (Supplementary Fig. 2).

**Supplementary Note 2** **Pipeline for haplotype-aware methylation calling using nanopore data**

The pipeline for haplotype-aware methylation calling using nanopore data is similar to the pipeline using PacBio data (Supplementary Fig. 3). In this pipeline, we used Guppy (version 4.2.2+effbaf8) to basecall nanopore raw reads. We then used Tombo^14^ (version 1.5.1) to re-squiggle the raw signals in nanopore reads to the reference genome, and then used DeepSignal2 (v0.1.2, https://github.com/PengNi/deepsignal2)^15^ to call 5mCpGs. We used Clair3^16^ (v0.1-r11 minor 2) with “*r941_prom_hac_g360+g422*” model to call variants. The called “PASS” SNVs were then used by WhatsHap^17^ (version 1.4) to assign the reads to two haplotypes. After generating phased methylation profiles by DeepSignal2, we used DSS (version 2.44.0)^13^ to get DMRs.

**Supplementary Note 3** **The model architecture of ccsmeth**

(1) bidirectional GRU

A bidirectional GRU^18^ layer includes a forward GRU and a backward GRU to catch both the forward and reverse flow of features. Suppose *x_1_, x_2_,…, x_t_* are a sequence of features, each time step $x_{i}$ contains four features: the nucleotide base, the mean IPD value, the mean PW value, and the number of subreads. A GRU cell will recursively calculate the hidden layer *h* as follows:

$r_{t}=sigmoid(W_{r}[h_{t-1}, x_{t}]+b_{r})$	(**2**)

$z_{t}=sigmoid(W_{z}[h_{t-1}, x_{t}]+b_{z})$	(**3**)

$\hat{h}_{t}=tanh(W_{h}\cdot[r_{t}h_{t-1}, x_{t}]+b_{h})$	(**4**)

$h_{t}=\left( 1-z_{t} \right)h_{t-1}+ z_{t}\hat{h}_{t}$	(**5**)

where *W* and *b* are weight matrices and biases. *x_t_* is the input feature; *r_t_* is a reset gate; *z_t_* is an update gate; *h_t_* is the hidden state; and $\hat{h}_{t}$ represents information that needs to be updated in the current cell. The outputs of forward and backward GRU are combined as:

$h_{t,A}=h_{t,F}\bigoplus h_{t,B}$	(**6**)

(2) Bahdanau attention

Bahdanau attention^19^ receives all the hidden states of RNN cells and outputs context vector $c_{t}$as follows:

$score\left( h_{t}, h_{s} \right)=tanh(W_{1}h_{t}+{W_{2}h}_{s})$	(**7**)

$a_{ts}=softmax(score\left( h_{t}, h_{s} \right))$	(**8**)

$c_{t}= a_{ts}{h_{t}}^{T}$	(**9**)

where *h_t_* represents the hidden state in the output vector of BiGRU; *h_s_* contains the final hidden state for an element in the sequence from GRU; *W_1_* and *W_2_* are weight matrices.

(3) Softmax activation function to output methylated/unmethylated probabilities

A softmax activation layer is used in ccsmeth to predict the methylated and unmethylated probabilities of one sample as follows:

$softmax\left( x_{i} \right)=\frac{e^{x_{i}}}{\sum_{j=0}^{1} e^{x_{j}}}, i=0 or 1$	(**10**)

where *x_0_* and *x_1_* are two outputs from the former fully connected layer, for calculating unmethylated and methylated probabilities, respectively.

(4) The cross-entropy loss function used for training the read-level model is as follows:

$L_{CE}=z*-log\left( y \right)+\left( 1-z \right)*-log(1-y)$	(**11**)

where *z* is the true label vector and *y* is the predicted methylated probability vector from the softmax function.

(5) The mean squared error (MSE) loss function for training the site-level model is as follows:

$L_{MSE}= \frac{\sum_{i=0}^{n} {{(x}_{i}}^{2}- {y_{i}}^{2})}{n}$	(**12**)

where *x_i_* is the predicted methylation frequency, *y_i_* is the true methylation frequency, *n* is the number of samples.

**Supplementary Note 4** **Testing ccsmethphase using CCS data of the HN0641 family trio**

We sequenced three human samples of a Chinese family trio using CCS, and got 2 SMRT cells of CCS reads for each of the three samples: HN0641_FA (father), HN0641_MO (mother), HN0641_S1 (son). We tested ccsmethphase using the CCS reads of the family trio. As shown in Supplementary Fig. 11a, the results indicate that the three samples have similar methylation levels. 11.9%, 12.0%, 10.4% CpGs of HN0641_FA, HN0641_MO, HN0641_S1, respectively, have low (≤0.3) methylation frequencies, while 75.4%, 73.4%, 80.6% CpGs of the three samples, respectively, have high (≥0.7) methylation frequencies.

We then examined the haplotype-aware methylation status of known imprinted regions in HN0641_S1. As shown in Supplementary Fig. 11b, the well-characterized imprinted intervals show large methylation differences between the two haplotypes (median=0.51). 14.0% of other known imprinted intervals also show large (>0.5) methylation differences. The result of HN0641_S1 is consistent with the results in HG002 and SD0651_P1.

Using ccsmethphase, we generated 2,813 DMRs from the CCS data of HN0641_S1. The DMRs cover 108 (52.9%) of the known imprinted intervals (*i.e.*, 108 known imprinted intervals are overlapped with the CCS-generated DMRs) (Supplementary Fig. 11c). Moreover, the haplotype phasing results of the family trio show that ccsmethphase not only can detect imprinted intervals, but also reveals the pattern of parental imprinting correctly. In the results of ccsmethphase, the maternally imprinted intervals (*e.g.*, *GNAS_Ex1A* and *PEG10*) in HN0641_S1 show high methylation levels in the haplotype inherited from mother, while the paternally imprinted intervals (e.g., *MEG3*) show high methylation levels in the haplotype inherited from father (Supplementary Figs. 12-14).

**Supplementary Note 5** **Running time of the ccsmethphase pipeline**

We evaluated the running time of 8 main processes in the ccsmethphase pipeline: *SAMTOOLS_index_bam* for indexing the CCS bam files, *CCSMETH_call_mods*	for calling methylation in CCS reads, *PBMM2* for aligning CCS reads to the reference genome, *SAMTOOLS_merge_bam* for merging alignment bam files of the same “sample”, *CLAIR3* for calling SNVs, *WHATSHAP_phase_haplotag* for phasing SNVs and reads, *CCSMETH_call_freq* for calling methylation frequencies of CpGs, and *DSS* for calling DMRs. The data used for evaluation include 10 SMRT cells of CCS reads used for testing in this study, in which there are 2 SMRT cells of CCS reads for each of the 5 “samples”: HG002 (15Kb), HG002 (20Kb), HG002 (24Kb), CHM13 (20Kb), and SD0651_P1 (15Kb) (Supplementary Table 2). The evaluation was performed at an HPC cluster containing two kinds of servers: (1) Server-CPU with 48 CPU cores (Intel(R) Xeon(R) Gold 6248R CPU @ 3.0GHz) and 192 GB RAM; (2) Server-GPU with 40 CPU cores (Intel(R) Xeon(R) Gold 6248 CPU @ 2.50GHz), 384 GB RAM, and 2 Nvidia Tesla V100 GPU cards. Details of the applied computing resources for the processes are shown in Supplementary Table 15. The running time of the processes is shown in Supplementary Fig. 15. Note that for the first 3 processes, the running time for each SMRT cell is shown. For the last 5 processes, the running time for each “sample” (2 SMRT cells) is shown. The evaluation indicates that for a human sample with 2 SMRT cells of CCS reads, methylation phasing and ASM detection can be performed in less than 14 hours using ccsmethphase even on a single server (Supplementary Fig. 15).
